# PacBio Hi-Fi genome assembly of the Iberian dolphin freshwater mussel *Unio delphinus* Spengler, 1793

**DOI:** 10.1101/2023.01.16.524251

**Authors:** Gomes-dos-Santos André, Lopes-Lima Manuel, Machado M. André, Teixeira Amílcar, C. Castro L. Filipe, Froufe Elsa

**Author notes:** corresponding author(s): André Gomes-dos-Santos, Manuel Lopes-Lima, Elsa Froufe.

## Abstract

Mussels of order Unionida are a group of strictly freshwater bivalves with nearly 1,000 described species widely dispersed across world freshwater ecosystems. They are highly threatened showing the highest record of extinction events within faunal taxa. Conservation is particularly concerning in species occurring in the Mediterranean biodiversity hotspot that are exposed to multiple anthropogenic threats, possibly acting in synergy. That is the case of the dolphin freshwater mussel *Unio delphinus* Spengler, 1793, endemic to the western Iberian Peninsula with recently strong population declines. To date, only four genome assemblies are available for the order Unionida and only one European species. We present the first genome assembly of *Unio delphinus*. We used the PacBio HiFi to generate a highly contiguous genome assembly. The assembly is 2.5 Gb long, possessing 1254 contigs with a contig N50 length of 10 Mbp. This is the most contiguous freshwater mussel genome assembly to date and is an essential resource for investigating the species’ biology and evolutionary history that ultimately will help to support conservation strategies.

## Background & Summary

The application of genomics approaches to study non-model organisms is deemed a key approach to assess biodiversity and guide conservation ^1–4^. Whole genome assemblies (WGS), provide access to a species’ “entire genetic code”, thus representing the most comprehensive framework to efficiently decipher a species’ biology ^5,6^. Genomic resources allow accurate definition of conservation units, identification of genetic elements with conservation relevance, inference of adaptive potential, assessment of population health, as well as provide predictive assessments of the impact of human-mediated threats and climate change ^3,5,7,8^. Consequently, WGS and other genomic tools are key resources to study and guide conservative actions and management planning.

Bivalves of the Order Unionida (known as freshwater mussels) are commonly found throughout most of the world’s freshwater ecosystems, where they play key ecological roles (e.g., nutrient and energy cycling and retention)^9–11^ and provide important services (e.g., water clearance, sediment mixing, pearls, and other raw materials)^9,10,12^. Despite their indisputable importance for freshwater ecosystems, freshwater mussels are among the most threatened taxa, with many populations worldwide having well-documented records of continuous declines over the last decades, as well as of many local and global extinctions ^13–16^. Threatened species with limited distributions, such as the dolphin freshwater mussel *U. delphinus* Spengler, 1793 (Unionida: Unionidae) only found in the western Iberian Peninsula region (Fig. 1), represent particularly urgent but challenging targets for conservation ^17^. The dolphin freshwater mussel, *U. delphinus*, only recently recognised as a valid species ^18^, has been strongly affected by a series of human-mediated actions over the last decades, including habitat destruction, dams or barrier construction, pollution, poor river management, water depletion, the introduction of invasive species, among others ^17,19^. All these pressures are further augmented by the effects of climate change, especially the steep volatility of water annual cycles, which is particularly evident in the Mediterranean region ^20,21^. As a consequence, the current area of occurrence of the dolphin freshwater mussel has been reduced by almost one-third from its historical distribution^19^. This concerning trend has triggered an unprecedented effort to research threats and promote and implement conservation policies. These are critically dependent on the understanding of the multiple aspects of the species’ biology, such as its life history, reproductive demands, ecological requirements, and its abiotic and biotic interactions ^13,17,19,22^.

**Fig. 1.**
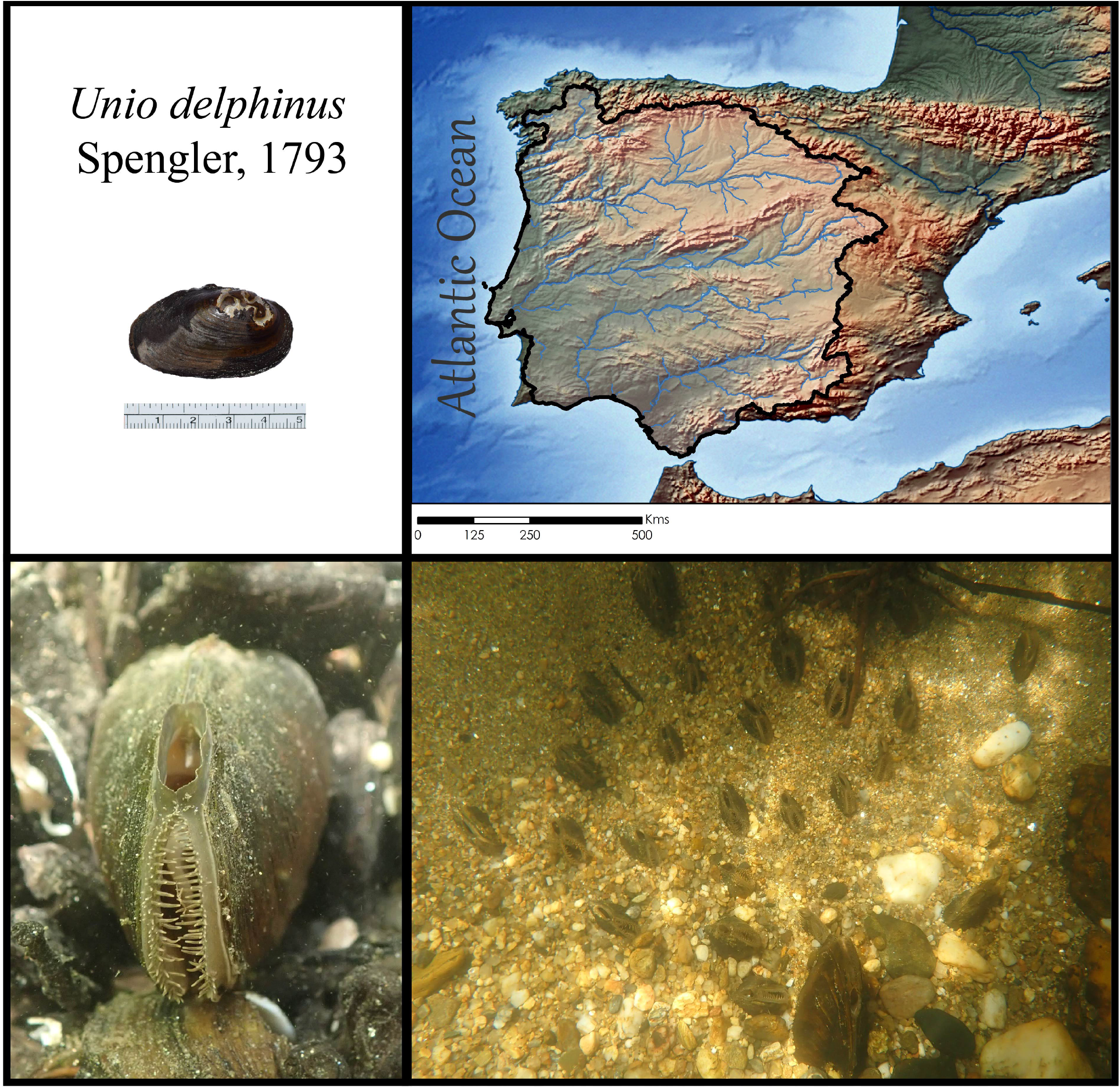
Top left: The *Unio delphinus* specimen used for the whole genome assembly. Top Right: The map of the *Unio delphinus* potential distribution produced by overlapping points of recent presence records (obtained from ^13^) with the Hydrobasins level 5 polygons ^68^. Bottom Left: An *Unio delphinus* individual in its natural habitat. Bottom Right: A population of *Unio delphinus* in their natural habitat (Photos by Manuel Lopes-Lima).

Recent efforts have focused on providing a thorough characterization of the species’ genetic diversity, population structure, and evolutionary history ^22–24^. Despite the unarguable accomplishments of these early molecular studies, the availability of large-scale and more biologically informative genomics resources is almost inexistent, not only for *U. delphinus* but also for all freshwater mussels. In fact, for approximately 1000 known species, only four whole genome assemblies ^25–28^ and less than 20 transcriptomes are currently available ^29–42^. Recently, the first transcriptome assemblies of five threatened European freshwater mussels species have been published, including the gill transcriptome of the dolphin freshwater mussel ^42^. This transcriptome represented a fundamental tool to start studying the evolutionary and adaptive traits of the species. However, single tissue RNA-seq approaches comprehend only a small fraction of the genetic information. Conversely, whole genome sequence assemblies represent a critically informative and research fertile resource to investigate and decipher multiple aspects of the species’ biology.

Here, we provide the first whole genome assembly of the dolphin freshwater mussel, *U. delphinus*. This represents the most contiguous freshwater genome assembly available, and the first Unionidae freshwater genome assembly from a European species. This genome represents a unique tool for an in-depth exploration of the many molecular mechanisms that govern this species’ biology, which will ultimately guide conservation genomic studies to protect the critical declining population trend.

## Methods

### Animal sampling

One individual of *Unio delphinus* was collected in the Rabaçal River in Portugal (Table 1) and transported alive to the laboratory, where tissues were separated, flash-frozen, and stored at −80⍰°C. The shell and tissues are deposited at CIIMAR tissue and mussels’ collection.

**Table 1.** MixS descriptors for the *Unio delphinus* specimen used for whole genome sequencing.

| **Sample** | ***Unio delphinus*** |
| --- | --- |
| Investigation_type | Eukaryote |
| Lat_lon | 41.564361; -7.258665 |
| Geo_loc_name | Portugal |
| Collection_date | 3/20/2021 |
| Env_package | Water |
| Collector | Amilcar Teixeira |
| Sex | Undetermined |
| Maturity | Mature |

### DNA extraction, library construction, and sequencing

For PacBio HiFi sequencing, mantle tissue was sent to Brigham Young University (BYU), where high-molecular-weight DNA extraction and PacBio HiFi library construction and sequencing were performed, following the manufacturer’s recommendations (https://www.pacb.com/wp-content/uploads/Procedure-Checklist-Preparing-HiFi-SMRTbell-Libraries-using-SMRTbell-Express-Template-Prep-Kit-2.0.pdf). Size-selection was conducted on the SageELF system. Sequencing was performed on four single-molecule, real-time (SMRT) cells using Sequel II system v.9.0, with a run time of 30 h, and 2.9 h pre-extension. The circular consensus analysis was performed in SMRT^®^ Link v9.0 (https://www.pacb.com/wp-content/uploads/SMRT_Link_Installation_v90.pdf) under default settings (Table 2).

**Table 2.**
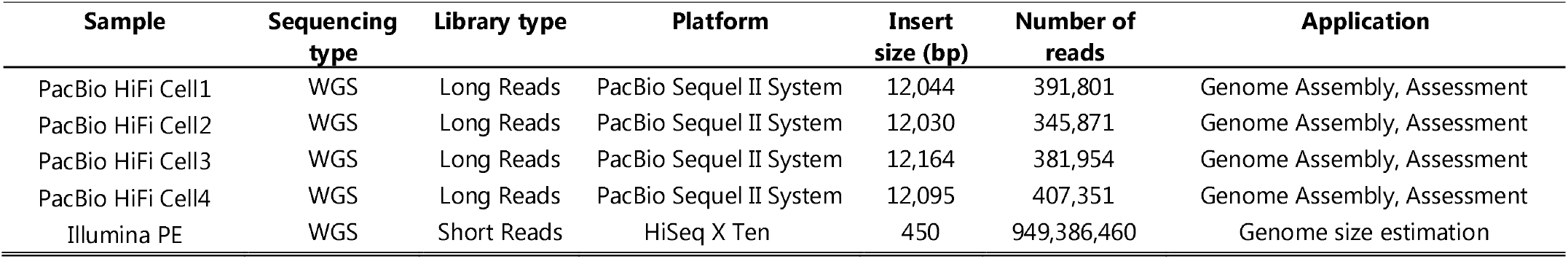
General statistics of raw sequencing reads used for the *Unio delphinus* genome assembly.

For short read Illumina sequencing, extracted genomic DNA was sent to Macrogen Inc. where a standard Illumina Truseq Nano DNA library preparation and whole genome sequencing of 150⍰bp paired-end (PE) reads was achieved using an Illumina HiSeq X machine (Table 2).

### Pre-assembly processing

Before the assembly, the characteristics of the genome were accessed with a k-mer frequency spectrum analysis using the PE reads. Briefly, read quality was evaluated using FastQC (https://www.bioinformatics.babraham.ac.uk/projects/fastqc/) and after, reads were quality trimmed with Trimmomatic v.0.38 ^43^, specifying the parameters “LEADING: 5 TRAILING: 5 SLIDINGWINDOW: 5:20 MINLEN: 36”. The quality of the clean reads was validated in FastQC. Genome size estimation was performed using the clean reads using Jellyfish v.2.2. and GenomeScope2 ^44^ specifying the k-mer length of 21.

### Mitochondrial genome assembly

PacBio HiFi reads were used to retrieve a whole mitochondrial genome (mtDNA) assembly by applying a pipeline recently developed by our group ^45^. Briefly, all Unionida mtDNA assemblies available on NCBI were retrieved (Fasta format; Retrieved in November 2022) and used as a reference mitogenome database. All the raw PacBio HiFi reads were mapped to the mitogenome database using Minimap2 v.2.17 ^46^, specifying parameters (-ax asm20). The output sam file was converted to bam and sorted using Samtools v.1.9^47^, with options “view” and “sort”, respectively. Samtools “view” was also used to retrieve only the mapped reads with parameter (-F 0×04) and after the bam file was converted to fastq format using the option “bam2fq”. The resulting PacBio HiFi mtDNA reads were corrected using Hifiasm v.0.13-r308 ^48,49^ with parameters (–write-ec). The corrected reads were assembled using Unicycler v.0.4.8 ^50^, a software package optimised for circular assemblies, with default parameters. Mitogenome annotation was produced using MitoZ v.3.4 ^51^ with parameters (--genetic_code 5 --clade Mollusca), using the PE reads for coverage plotting.

### Genome assembly

The overall pipeline used to obtain the genome assembly and annotation is provided in Fig. 2.

**Fig. 2.**
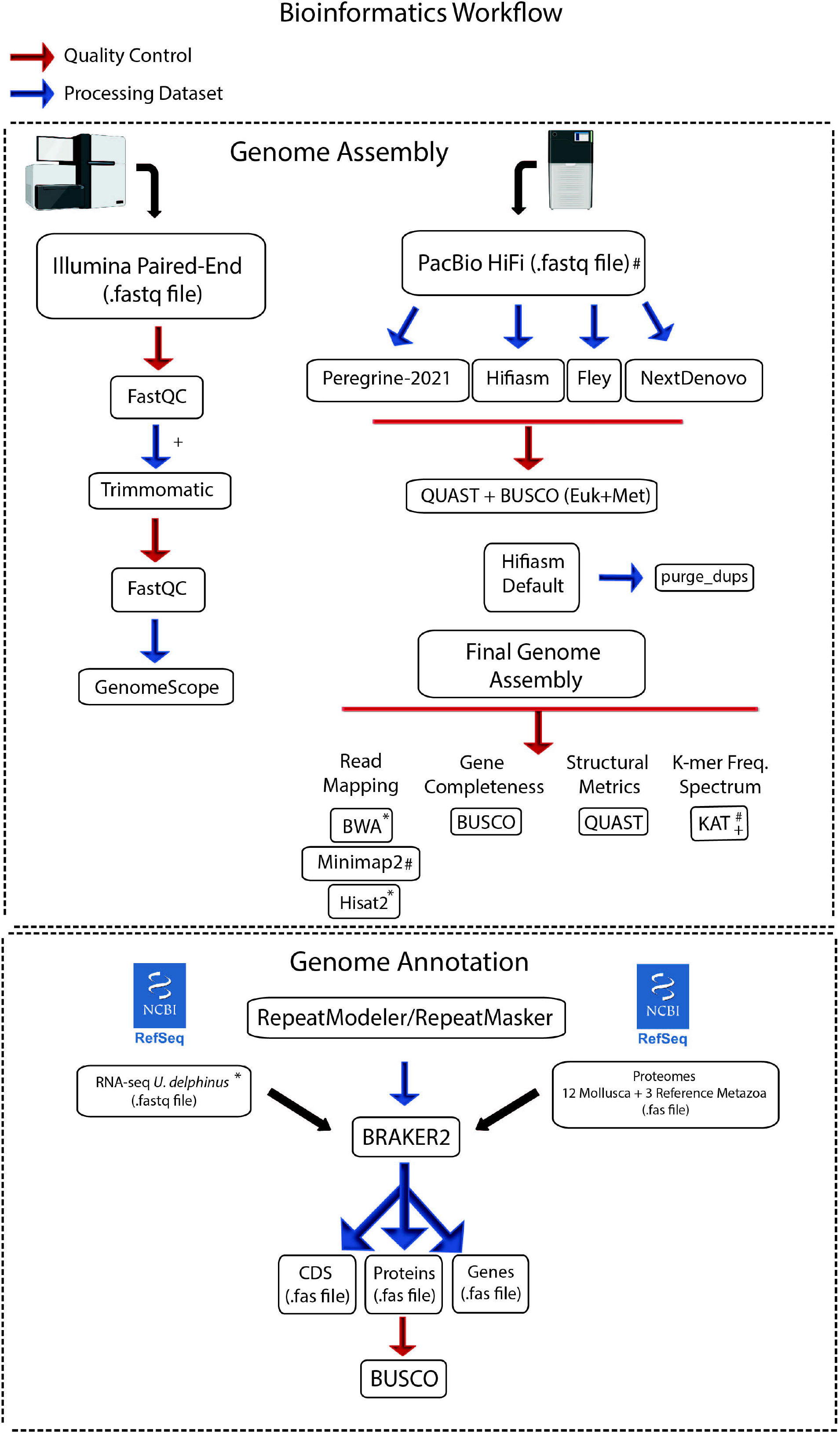
Bioinformatics pipeline applied for the whole genome assembly and annotation. Representative figures created with BioRender.com.

Firstly, PacBio HiFi reads were assembled using multiple software optimized for PacBio HiFi reads, i.e., Hifiasm 0.16.1-r375^48,49^ with default parameters, Flye v.2.8.3^52^ with parameters (--pacbio-hifi), NextDenovo v.2.4.0 (https://github.com/Nextomics/NextDenovo) with parameters (read_type = hifi) and peregrine-2021 v0.4.3^53^ with default parameters. After, the overall quality of each assembly was accessed using Benchmarking Universal Single-Copy Orthologs (BUSCO) v.5.2.2^54^ with Eukaryota and Metazoa databases and Quality Assessment Tool for Genome Assemblies (QUAST) v.5.0.2^55^ (Fig. 2). Hifiasm v.0.13-r308 produced the best results of the tested assemblies and thus was selected for further analyses. Since the genome size was larger than predicted by the GenomeScope report, several new assemblies were produced with this Hifiasm v.0.13-r308, testing a range of parameters (*l* = 3; s = 0.50, 0.45, 0.35), following the authors’ recommendations (https://hifiasm.readthedocs.io/en/latest/faq.html#p-large). Given that reducing the similarity threshold for duplicate haplotigs (i.e., parameter -l and -s) had little impact on the overall statistic, the assembly with default parameters was chosen for further analysis. To separate poorly resolved pseudo-haplotypes, purge_dups v.1.2.5^56^ was applied, first with default parameters and after by manually adjusting the transition between haploid and diploid cutoff (i.e., parameter -m of option calcuts) to 30, 32 and 25 in three independent runs. In all the runs the lower and upper bound for read depth were always maintained, i.e., 5 and 87, respectively. All the cutoff values were determined by inspection of the *k*-mer plot produced by the K-mer Analysis Toolkit (KAT) tool^57^. The influence of purge_dups v.1.2.5 was evaluated using BUSCO v.5.2.2 with Eukaryota and Metazoa databases and QUAST v.5.0.2. Since purge_dups v.1.2.5 did not remove any duplicates (neither with the default nor adjusted cutoffs) the Hifiasm 0.16.1-r375 default assembly was selected as the final assembly. To evaluate the quality of the final assembly, several metrics and software were used. Besides BUSCO v.5.2.2 and QUAST v.5.0.2 metrics, completeness, heterozygosity, and collapsing of repetitive regions were evaluated using a k-mer distribution with KAT ^57^. Moreover, read-back mapping was performed for the PE using with Burrows-Wheeler Aligner (BWA) v.0.7.17-r1198^58^, for long reads with Minimap2 v.2.17 and for RNA-seq (SRR19261764, ^42^) with Hisat2 v.2.2.0^59^.

### Masking of repetitive elements, gene models predictions and annotation

Before masking repetitive elements, a *de novo* library of repeats was created for the final genome assembly, with RepeatModeler v.2.0.133^60^. Subsequently, the genome was soft masked combining the *de novo* library with the ‘Bivalvia’ libraries from Dfam_consensus-20170127 and RepBase-20181026, using RepeatMasker v.4.0.734 ^61^.

The masked assembly was used for gene prediction, performed using BRAKER2 pipeline v2.1.6 ^62^. First, RNA-seq data from *U. delphinus* was retrieved from GenBank (SRR19261764, ^42^) (the same individual used for the genome assembly), quality trimmed with Trimmomatic v.0.3839 (parameters described above) and aligned to the masked genome, using Hisat2 v.2.2.0 with the default parameters. Moreover, the complete proteome of 14 mollusc species and three reference Metazoa genomes (*Homo sapiens, Ciona intestinalis, Strongylocentrotus purpuratus*), were used as supplementary evidence for gene prediction, were downloaded from public databases (Table 3). BRAKER2 pipeline was applied, specifying parameters “-etpmode; –softmasking;” and subsequently, gene predictions were renamed, cleaned, and filtered using AGAT v.0.8.0^63^, which also corrected overlapping prediction, removed coding sequence regions (CDS) with <100 amino acid and removed incomplete gene predictions (i.e., without start and/or stop codons). Finally, proteins extracted with AGAT were used for functional annotation, using InterProScan v.5.44.80 ^64^ and BLASTP searches against the RefSeq database^65^. Homology searches were obtained using DIAMOND v.2.0.11.149 ^66^, specifying the parameters “-k 1, -b 20, -e 1e-5, --sensitive, --outfmt 6”. Finally, BUSCO scores were estimated for the predicted proteins, using the Eukaryota and Metazoa databases, as described above.

**Table 3.**
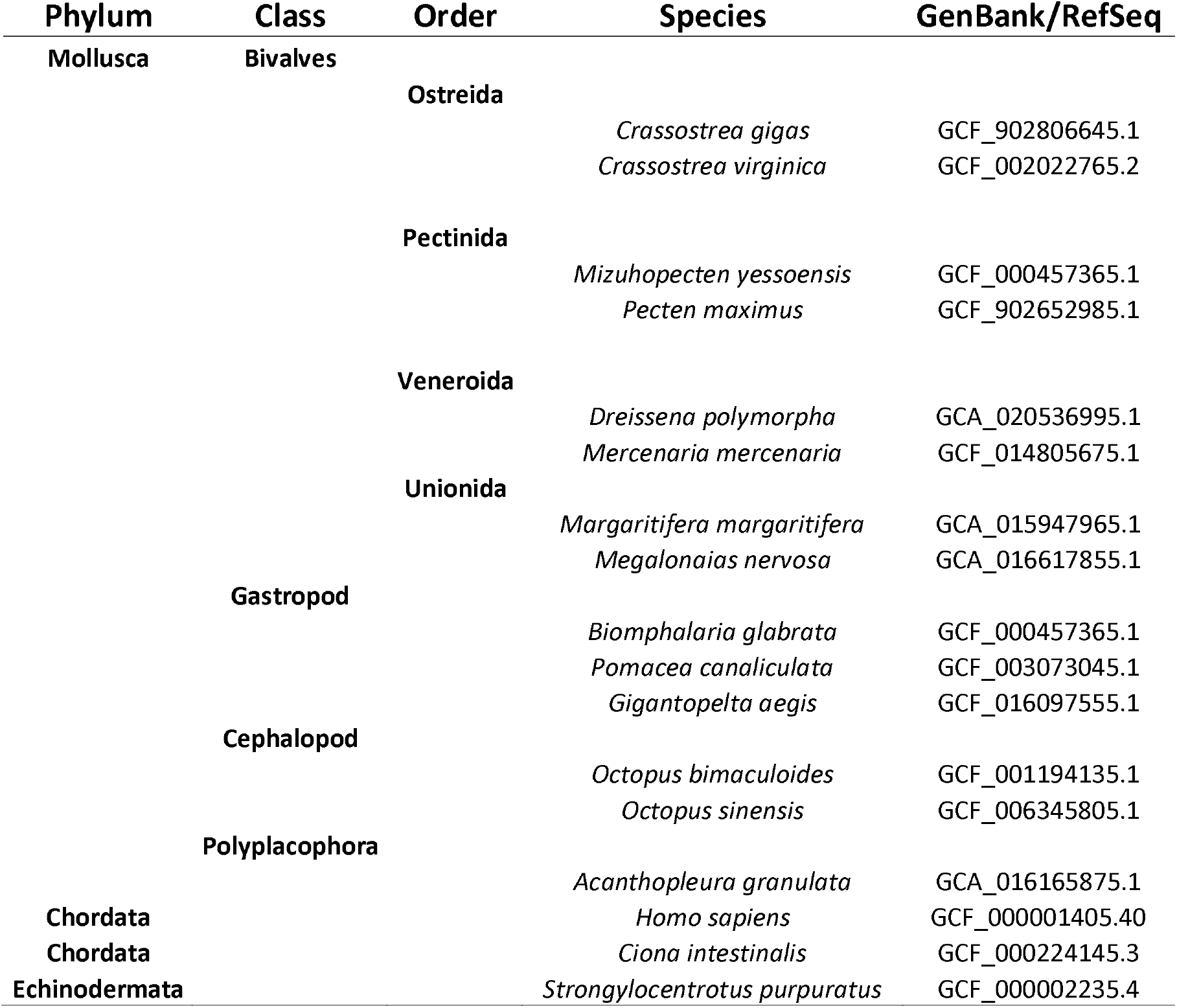
List of proteomes used for BRAKER2 gene prediction pipeline.

## Data Records

The raw reads sequencing outputs were deposited at the NCBI Sequence Read Archive with the accession’s numbers: SRR23060683, SRR23060685, SRR23060678 and SRR23060675 for PacBio CCS HiFi; SRR23060686 for Illumina PE. The Genome assembly is available under accession number JAQISU000000000. BioSample accession number is SAMN32554582 and BioProject PRJNA917855. The remaining information was uploaded to figshare (10.6084/m9.figshare.21878946). In detail, the files uploaded to figshare include the final unmasked and masked genome assemblies (Ude_BIV7592_haploid.fa and Ude_BIV7592_haploid_SM.fa), the annotation file (Ude_BIV7592_annotation_v4.gff3), predicted genes (Ude_BIV7592_genes_v4.fasta), predicted messenger RNA (Ude_BIV7592_mrna_v4.fasta), predicted open reading frames (Ude_BIV7592_cds_v4.fasta), predicted proteins (Ude_BIV7592_proteins_v4.fasta), as well as full table reports for Braker gene predictions and InterProScan functional annotations (Ude_BIV7592_annotation_v4_InterPro_report.txt) and RepeatMasker predictions (Ude_BIV7592_annotation_v4_RepeatMasker.tbl).

## Technical Validation

### Raw datasets and pre-assembly processing quality control

Raw sequencing outputs general statistics are provided in Table 2. GenomeScope2 estimated genome size was ^~^1.31 Gb and heterozygosity levels of ^~^46.8% (Fig. 3a), which are within the values observed for other Unionidae genomes available ^25–28^.

**Fig. 3.**
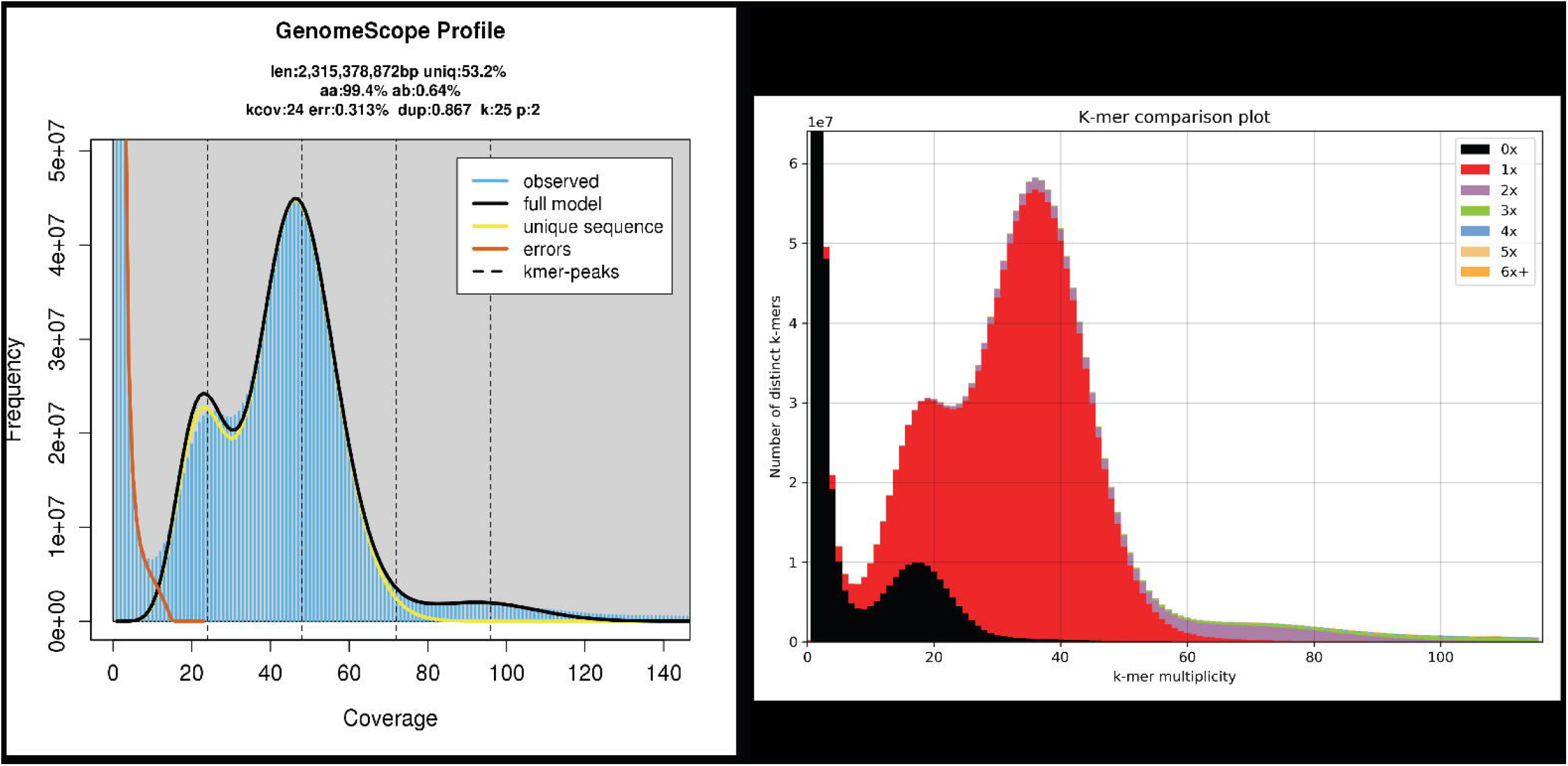
Left: GenomeScope2 k-mer (21) distribution displaying estimation of genome size (len), homozygosity (aa), heterozygosity (ab), mean k-mer coverage for heterozygous bases (kcov), read error rate (err), the average rate of read duplications (dup), k-mer size used on the run (k:), and ploidy (p:). Right: *Unio delphinus* genome assembly assessment using KAT comp tool to compare the PacBio HiFi k-mer content within the genome assembly. Different colours represent the read k-mer frequency in the assembly.

### Genome assembly metrics

Hifiasm produced the overall best genome assembly of all the tested assemblers (Table 4). Both Fley and peregrine-2021 were very inefficient in collapsing haplotypes, resulting in unexpectedly large assemblies with high levels of duplicated BUSCO scores (Table 4). Conversely, Hifiasm and NextDenovo efficiently resolve duplicates while ensuring high complete BUSCO scores (Table 4). Additionally, Hifiasm produced a much more contiguous genome assembly, with an almost 5-fold increased N50 length (Tables 3–4). Although the BUSCO scores of the Hifiasm assembly had residual percentages of duplicated sequences, considering the increased genome size compared with GenomeScope estimation, as well as the genome sizes of other Unionidae assemblies (Table 5), we tested several similarity thresholds for duplicates in Hifiasm. The impact of the resulting assemblies on the overall statistics was limited, i.e., -s 0.50-0.35, or had no impact at all, i.e., -l 3 (Table 5). Although two of the assemblies, i.e., -s 0.50 and -s 0.45, show a slight increase in the N50 length (Table 5), given the overall little impact in the final genome size, we opted to use the Hifiasm default assembly as the final assembly. Moreover, purg-dups software did not remove any additional sequences from the Hifiasm default assembly, suggesting that reducing the similarity threshold for duplicate haplotigs (option -s) might be over-purging the assembly.

**Table 4.**
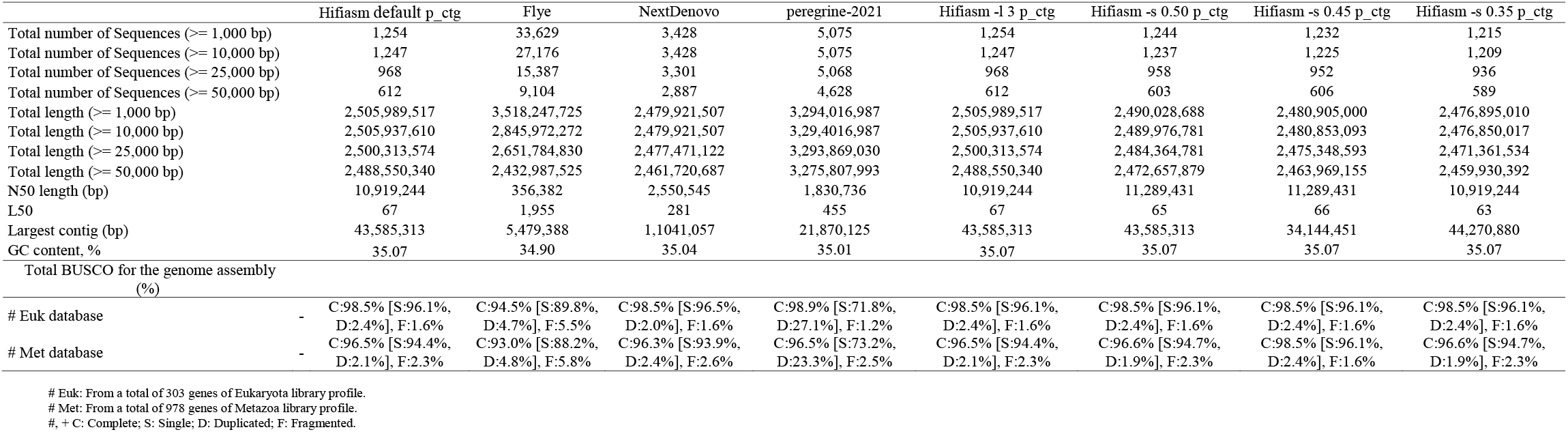
*Unio delphinus* genome assemblies tests’ general statistics.

**Table 5.** General statistics of the *Unio delphinus* final genome assembly (p_ctg); *Unio delphinus* alternative haplotypes genome assemblies (hap1 and hap2); other published freshwater mussels genome assemblies.

|  |  | Hifiasm -l 3 p_ctg | Hifiasm -l 3 hap1 | Hifiasm -l 3 hap2 | Megaloniais nervosa | Potamilus streckeri | Margaritifera margaritifera |
| --- | --- | --- | --- | --- | --- | --- | --- |
| Total number of Sequences (>= 1,000 bp) |  | 1,254 | 3,752 | 3,000 | 90,895 | 2,366 | 105,185 |
| Total number of Sequences (>= 10,000 bp) |  | 1,247 | 3,743 | 2,993 | 54,764 | 2,162 | 15,384 |
| Total number of Sequences (>= 25,000 bp) |  | 968 | 2,774 | 2,668 | 29,042 | 1,831 | 11,583 |
| Total number of Sequences (>= 50,000 bp) |  | 612 | 1,938 | 2,029 | 12,699 | 1,641 | 9,265 |
| Total length (>= 1,000 bp) |  | 2,505,989,517 | 2,311,195,669 | 2,291,510,236 | 2,361,438,834 | 1,776,751,942 | 2,472,078,101 |
| Total length (>= 10,000 bp) |  | 2,505,937,610 | 2,311,130,750 | 2,291,456,057 | 2,193,448,794 | 1,775,453,721 | 2,293,496,118 |
| Total length (>= 25,000 bp) |  | 2,500,313,574 | 2,293,207,905 | 2,285,083,051 | 1,768,523,103 | 1,769,874,087 | 2,236,013,546 |
| Total length (>= 50,000 bp) |  | 2,488,550,340 | 2,264,885,011 | 2,262,774,153 | 1,194,323,847 | 1,763,052,140 | 2,152,307,394 |
| N50 length (bp) |  | 10,919,244 | 4,974,507 | 4,544,314 | 50,662 | 2,051,244 | 288,726 |
| L50 |  | 67 | 125 | 121 | 12,463 | 245 | 2,393 |
| Largest contig (bp) |  | 43,585,313 | 27,621,201 | 28,529,984 | 588,638 | 10,787,299 | 2,510,869 |
| GC content, % |  | 35.07 | 35.07 | 35.04 | 35.82 | 33.79 | 35.42 |
| Clean Paired-End (PE) Reads Alignment Stats |  |  |  |  |  |  |  |
| Percentage of Mapped WGS PE (%) | - | 99.81% | - | - | - | - | - |
| Percentage of Mapped WGS PacBio (%) |  |  | - | - | - | - | - |
| Percentage of Mapped RNA-seq PE (%) |  | 96.15% | - | - | - | - | - |
| Total BUSCO for the genome assembly (%) |  |  |  |  |  |  |  |
| # Euk database | - | C:98.5% [S:96.1%, D:2.4%], F:1.6% | C:94.2% [S:91.8%, D:2.4%], F:3.5% | C:92.9% [S:90.2%, D:2.7%], F:3.1% | C:70.6% [S:70.2%, D:0.4%], F:14.9% | C:98.1% [S:97.3%, D:0.8%], F:0.8% | C: 86.8% [S: 85.8%, D:1.0%], F: 5.9% |
| # Met database | - | C:96.5% [S:94.4%, D:2.1%], F:2.3% | C:92.1% [S:90.5%, D:1.6%], F:3.5% | C:92.1% [S:90.4%, D:1.7%], F:3.5% | C:71.5% [S:70.1%, D:1.4%], F:14.5% | C:95.0% [S:93.6%, D:1.4%], F:2.3% | C: 84.9% (S: 83.8%, D: 1.1%), F: 4.9% |
| Masking Repetitive Regions and Gene Prediction |  |  |  |  |  |  |  |
| Percentage masked bases (%) | - | 52.83 | - | - | 25.00 | 51.03 | 59.07 |
| Number of mRNA | - | 44,382 | - | - | 49,149 | 41,065 | 40,544 |
| Protein coding genes (CDS) | - | 44,382 | - | - | 49,149 | 41,065 | 35,119 |
| Functional annotated genes |  | 32,089 | - | - | - | - | 31,584 |
| Total gene length (bp) | - | 869,540,056 | - | - | - | - | 902,994,752 |
| Total BUSCO for the predicted proteins (%) |  |  |  |  |  |  |  |
| + Euk database | - | C:96.8% [S:88.2%, D:8.6%], F:2.7% | - | - | - | - | C:90.6% [S:81.2%, D:9.4%], F:3.9% |
| + Met database | - | C:97.3% [S:86.0%, D:11.3%], F:2.3% | - | - | - | - | C:92.6% [S:82.3%, D:10.3%] , F:3.2% |
| # Euk: From a total of 303 genes of Eukaryota library profile.<br># Met: From a total of 978 genes of Metazoa library profile.<br>+ Euk: From a total of 255 genes of Eukaryota library profile.<br>+ Met: From a total of 954 genes of Metazoa library profile.<br>#, + C: Complete; S: Single; D: Duplicated; F: Fragmented. |  |  |  |  |  |  |  |

The final genome assembly has a total length of ^~^2.5 Gbp, which is relatively larger than the GenomeScope size estimation, i.e., ^~^2.31 Gbp (Table 5, Fig. 3a). Although unexpected, the fact is that from all the primary assemblies here produced (from different software and Hifiasm parameters), none had a total length close to those estimated from GenomeScope (Tables 4–5). The alternative haplotypes assemblies produced by Hifiasm have a total length similar to the GenomeScope estimations, however, the complete BUSCO scores were considerably reduced for these assemblies with no significate impact on duplicates (Table 5). On the other hand, purge-dups did not report any duplicated sequences in the assembly, which further support that Hifiasm efficiently resolved the haplotype variants. Moreover, the few genome assemblies available for freshwater mussels, show considerable distinct genome sizes (up to 696Mbp difference in size), even within the family Unionidae (Table 5). Consequently, the discrepancies between GenomeScope and the final genome size are likely a consequence of short read-based k-mer frequency spectrum analyses inaccurate estimation of the genome size.

The assembly here presented also shows, the most contiguous freshwater mussel genome assembly available to date, with a contig N50 length of ^~^ 10 Mbp, which represents a ^~^5-fold increase in N50 length regarding the only other PacBio-based genome assembly, i.e., from *P. streckersoni* ^26^ (Table 5). The levels of completeness reported by BUSCOs scores are also within those observed for other freshwater mussel genome assemblies, with nearly no fragmented nor missing hits for both the eukaryotic and metazoan curated lists of near-universal singlecopy orthologous (Table 5). The KAT k-mer analyses revealed a low level of k-mer duplication (blue, green, purple, and orange in Fig. 3b), with a high level of haplotype uniqueness (red in Fig. 3c) and a similar k-mer distribution to GenomeScope2 (performed with Illumina PE reads Fig. 3 a,b). Both short-read, RNA-seq and long-read back-mapping percentages resulted in an almost complete mapping (Table 5). Overall, these general statistics validate the high completeness, low redundancy, and quality of the final genome assembly.

### Repeat masking, gene models prediction, and annotation

RepeatModeler/RepeatMasker masked 52.83% of the genome, a value within those observed for other Unionida genome assemblies and close to the GenomeScope estimation (Table 6, Fig. 3a). Unlike the results observed in previous freshwater mussel’s genome assemblies ^25,26^, most repeats are classified as DNA elements (21.92%, ^~^ 549 Mgp), rather than unclassified (16.32 %, ^~^ 408 Mgp), with the remaining categories having similar percentages (Table 6). These results might be a consequence of PacBio HiFi reads efficiency in resolving repetitive regions thus facilitating their classification. BRAKER2 gene prediction identified 44,382 CDS, which is close to the predictions of the other freshwater mussel assemblies (Table 5). BUSCO scores for protein predictions showed almost no missing hits for either of the near-universal single-copy orthologous databases used (Table 5). The number of functionally annotated genes was 32,089, which is similar to the number of annotated genes for the *Margaritifera margaritifera* genome assembly (Table 5)^25^. Overall, the numbers of both predicted and annotated genes are within the expected range for bivalves (reviewed in ^67^), as well as within the records of other freshwater mussel assemblies (Table 5)^25–28^.

**Table 6.**
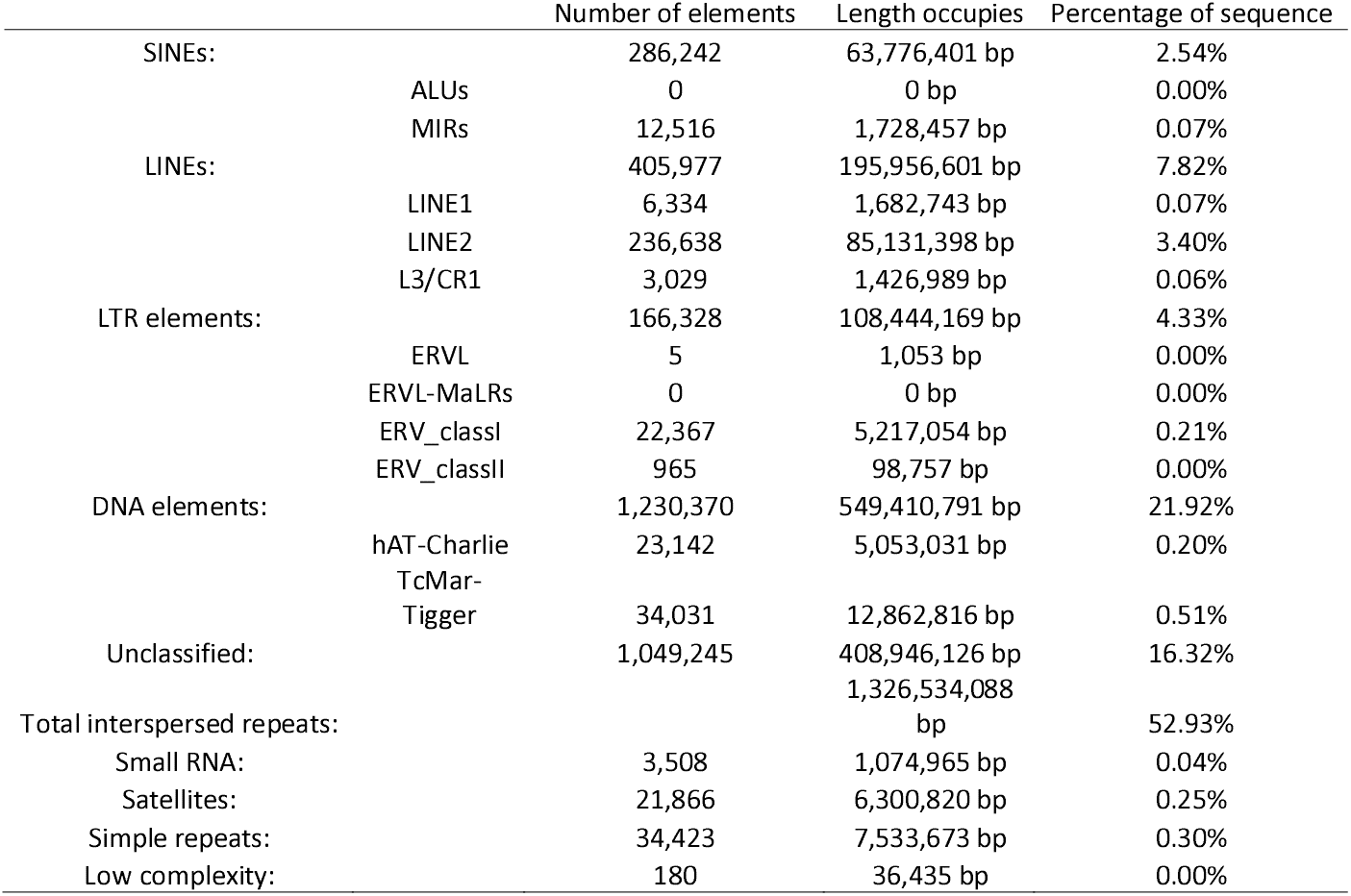
RepeatMasker report of the content of repetitive elements in the *Unio delphinus* genome assembly.

The results here presented revealed the significant impact that using PacBio HiFi long-read sequencing has on assembling freshwater mussels’ genomes, by producing the most contiguous freshwater mussel genome assembly to date. Furthermore, the overall quality and completeness of the genome are demonstrated using several distinct statistics and comparative approaches. This genome represents therefore a key resource to start exploring the many biological, ecological, and evolutionary features of this highly threatened group of organisms, for which the availability of genomic resources still falls far behind other molluscs.

## Code Availability

All software with respective versions and parameters used for producing the resources here presented (i.e., transcriptome assembly, pre and post-assembly processing stages, and transcriptome annotation) are listed in the methods section. Software programs with no parameters associated were used with the default settings.

## Acknowledgements

AGS was funded by the Portuguese Foundation for Science and Technology (FCT) under the grants SFRH/BD/137935/2018 and COVID/DB/152933/2022, which also supported MLL (2020.03608.CEECIND) and EF (CEECINST/00027/2021). This research was developed under the project EdgeOmics - Freshwater Bivalves at the edge: Adaptation genomics under climate-change scenarios (PTDC/CTA-AMB/3065/2020) funded by FCT through national funds. Additional strategic funding was provided by FCT UIDB/04423/2020 and UIDP/04423/2020.

## Author contributions

E.F, M.L.L, L.F.C.C designed and conceived this work.

M.L.L., and A.T. collected the samples.

A.G.S and A.M.M carry on all the analysis.

A. G. S. and E. F wrote the first version of the manuscript.

All authors read, revised, and approved the final manuscript.

